# Social isolation is linked to classical risk factors of Alzheimer’s disease-related dementias

**DOI:** 10.1101/2021.09.13.460121

**Authors:** Kimia Shafighi, Sylvia Villeneuve, Pedro Rosa-Neto, AmanPreet Badhwar, Judes Poirier, Vaibhav Sharma, Yasser Iturria-Medina, Patricia P Silveira, Laurette Dube, David Glahn, Danilo Bzdok

**Affiliations:** Department of Biomedical Engineering, Faculty of Medicine, McGill University, Montreal, Canada; Neurology and Neurosurgery Department, Montreal Neurological Institute (MNI), Faculty of Medicine, McGill University, Montreal, Canada; McConnell Brain Imaging Centre (BIC), MNI, Faculty of Medicine, McGill University, Montreal, Canada; Department of Psychiatry, McGill University, H3A 1A1, Montreal, Quebec, Canada; Douglas Mental Health University Institute, Studies on Prevention of Alzheimer’s Disease (StoP-AD) Centre, H4H 1R3, Montreal, Quebec, Canada; Translational Neuroimaging Laboratory, McGill University Research Centre for Studies in Aging, Alzheimer’s Disease Research Unit, Montreal, Canada; Le Centre intégré universitaire de santé et de services sociaux (CIUSSS) de l’Ouest-de-l’Île-de-Montréal, Montréal, Canada; Department of Neurology and Neurosurgery, Psychiatry and Pharmacology and Therapeutics, McGill University, Montreal, Canada; Département de pharmacologie et physiologie, Faculté de médecine, Université de Montréal, Montréal, Canada; Centre de recherche de l’Institut universitaire de gériatrie de Montréal (CRIUGM), Montréal, Canada; Centre for Studies in the Prevention of Alzheimer’s Disease, Douglas Mental Health Institute, McGill University, Montreal, Canada; Ludmer Centre for Neuroinformatics and Mental Health, McGill University, Montreal, Canada; Desautels Faculty of Management, McGill Center for the Convergence of Health and Economics, McGill University, Montreal, QC, Canada; Tommy Fuss Center for Neuropsychiatric Disease Research, Department of Psychiatry, Boston Children’s Hospital, Boston, MA, USA; Department of Psychiatry, Harvard Medical School, Boston, MA, USA; School of Computer Science, McGill University, Montreal, Canada; Mila - Quebec Artificial Intelligence Institute, Montreal, Canada

## Abstract

Alzheimer’s disease and related dementias is a major public health burden – compounding over upcoming years due to longevity. Recently, clinical evidence hinted at the experience of social isolation in expediting dementia onset. In 502,506 UK Biobank participants and 30,097 participants from the Canadian Longitudinal Study of Aging, we revisited traditional risk factors for developing dementia in the context of loneliness and lacking social support. Across these measures of subjective and objective social deprivation, we have identified strong links between individuals’ social capital and various indicators of Alzheimer’s disease and related dementias risk, which replicated across both population cohorts. The quality and quantity of daily social encounters had deep connections with key aetiopathological factors, which represent 1) personal habits and lifestyle factors, 2) physical health, 3) mental health, and 4) societal and external factors. Our population-scale assessment suggest that social lifestyle determinants are linked to most neurodegeneration risk factors, highlighting them promising targets for preventive clinical action.

## Introduction

Alzheimer’s disease and related dementias (ADRD) is a growing public health crisis. With no known cure, this devastating condition generates ~1 trillion global costs every year and places a considerable burden on patients, caregivers, and society.^1^ The number of ADRD cases is estimated to triple by 2050.^2^ In a parallel development, there is now rapidly growing evidence that social isolation is associated with an escalated risk of ADRD.^3–8^ In fact, the World Health Organization (WHO) has identified ADRD and social isolation, separately, as two global public health priorities.^9, 10^ Both challenges may now be aggravating due to social deprivation as a consequence of the COVID19 pandemic: many cities, states, and nations have imposed stringent social distancing measures – leading to probably the largest mass social isolation in recorded history.

Substantial progress has been made in delineating aetiopathological antecedents of this major neurodegenerative disease. While we have identified some biomarkers and short-term treatment of symptoms, our ability to attenuate the trajectory of neurodegenerative progression remains limited. As a source of hope, a recent consensus article^11^ reported that potentially modifiable factors in ADRD amount to as much as 40% of the overall disease risk. Widely agreed upon risk factors include childhood education, exercise, socioeconomic status, smoking, alcohol consumption, hearing and vision loss, depression, diabetes, hypertension, sleep apnea, air pollution and obesity.^11^ However, we still have a clouded understanding of how these risk factors are linked to social lifestyle. The relevance of subjective and objective social isolation for ADRD risk in relation to other commonly studied risk factors is only now attracting the attention of researchers, stakeholders, and policy makers. The premise of our study is that a wider characterization of these social behaviors in late life will enable a more complete conceptualization of ADRD risk, potentially paving the way for novel treatment avenues. Such new insight is imperative given that social behaviors are modifiable in principle through societal measures^12^ in contrast to genetically determined risk. This knowledge gap is particularly blatant when one considers social deprivation in the elderly. Social factors like loneliness, as a measure of *subjective* social isolation, and regular social support, as a measure of *objective* social isolation, are rarely considered in risk models or authoritative surveys of ADRD aetiopathology.

There is substantial evidence that acceleration in cognitive decline^13–15^ and increased dementia risk^16, 17^ co-occurs with loneliness in individuals, which is also indicated by greater ADRD-related neuropathological.^13, 18^ These pointers suggest that perceived social isolation plays an important and potentially independent role from objective social isolation in normative brain aging and its aberrations in neurodegenerative disease. Different facets of social isolation – loneliness, social network, social engagement, and social support – have been associated with poor health outcomes, including hypertension and immune system dysfunction,^19, 20^ cognitive decline,^14, 21, 22^ psychological distress (e.g. depression, anxiety), increased dementia risk^5^ and shortened life expectancy.^23^ Studying the role of social lifestyle in ADRD onset should therefore acknowledge determinants of both subjective loneliness feelings and objective social support frequency. Here, we have systematically revisited classical, widely acknowledged aetiopathological factors closely linked to ADRD by capitalizing on two unique cohorts: 502,506 participants from the UK Biobank^24^ and 30,097 participants from the Canadian Longitudinal Study of Aging.^25^ Empowered by the advent of the UK Biobank and CLSA cohorts,^26, 27^ we have tested the hypothesis that subjective loneliness and objective social support show robust associations with major ADRD risk factors. Narrowing this knowledge gap is particularly urgent when considering less well studied risk factors like late-life behaviors, including subjective and objective social isolation – which were recently exacerbated as a consequence of the COVID pandemic.

## Methods

### Population cohort 1: United Kingdom Biobank

The UK Biobank is a prospective epidemiological cohort that offers extensive behavioral and demographic assessments, medical and cognitive measures, as well as biological samples in 502,506 participants, which were recruited from across Great Britain.^28^ Our study involved the full population sample including 54.4% females, aged 40-69 years when recruited (mean age 56.5, standard deviation (SD) 8.1 years). The present analyses were conducted under UK Biobank application number 25163. All participants provided written, informed consent and the study was approved by the Research Ethics Committee (REC number 11/NW/0382). Further information on the consent procedure can be found elsewhere (http://biobank.ctsu.ox.ac.uk/crystal/field.cgi?id=200).

### Population cohort 2: Canadian Longitudinal Study of Aging

CLSA has been launched in 2011 as an independent prospective epidemiological cohort, which places a focus on aging trajectories and deep phenotyping.^29^ This study follows a population of 30,097 individuals, including 50.9% females, aged 44-89 at enrollment (mean age 63.0, SD 10.3 years), recruited from 11 cities in 10 provinces across Canada. The acquisition of baseline data finished in 2015. Ethics approval was obtained by the Research Ethics Board at McGill University (REB file #20-05-068).

### Data availability

UK Biobank data are available through a procedure described at http://www.ukbiobank.ac.uk/using-the-resource/. All CLSA data are accessible to researchers through data access requests at https://clsa-elcv.ca/data-access).

### Social isolation target phenotypes

Studying the role of social lifestyle in ADRD onset should acknowledge determinants of both subjective loneliness feelings and objective social support frequency. Regarding the loneliness target, we used the yes/no answer from UK Biobank participants to the question ‘Do you often feel lonely?’ (data field 2020). In CLSA, our loneliness target measure was based on the question ‘How often did you feel lonely?’, with the positive answer denoting ‘all of the time (5-7 days)’. The need for brief loneliness assessments has long been recognized, particularly for inclusion in large population-based studies ^30^. Regarding the social support target, our UK Biobank analyses were based on the question ‘How often are you able to confide in someone close to you?’, as an objective measure of the frequency of social interactions (data field 2110). Our study modeled lack of social support as less than ‘daily or almost daily’ (positive answer) against confiding in others more often (treated as negative answer). In CLSA, regarding lack of social support, participants were asked the question ‘Someone to confide in or talk to about yourself or your problems?’, where less than ‘all of the time’ or ‘most of the time’ was modeled as the positive case.

### Multivariate decomposition approach

We used partial least squares (PLS) correlation to examine possible cross-associations between classical ADRD risk factors and social richness indicators (cf. Supplementary Table 1). As used in our previous work, this technique is particularly useful when handling very large and strongly correlated datasets.^31^ To analyze the relationship between the risk traits and the social factors, the observations (here, the participant responses) were stored in matrices denoted by *X* and *Y*, respectively. The two sets of linear combinations of the original variables are obtained as follows:

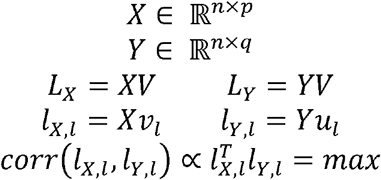

where *n* denotes the number of observations or individuals, *p* is the number of risk traits, *q* is the number of social factors (7 in the UKBB and 6 in the CLSA), V and U denote the respective contributions of *X* and *Y, L_X_* and *L_Y_* denote the respective latent ‘modes’ of joint variation between patterns in *X* and patterns in *Y, l_x,l_* is the *i*^th^ column of *L_X_*, and *l_Y,l_* is the *l*^th^ column of *L_Y_*. The goal of our PLS correlation application was to find pairs of latent vectors *l_X,l_* and *l_Y,l_* with maximal correlation in the derived latent embedding. Since PLS correlation was purely used as an explanatory analysis, uncertainty in effect sizes were not measured.

### Bayesian regression approach

Next, to ascertain robust associations between social richness and ADRD risk in the wider society, Bayesian hierarchical regression was a natural choice of method,^32^ following our previous work at the population level.^31, 33, 34^ In particular, classical tests for statistical significance would only have provided dichotomic statements in form of p-values against the null hypothesis of no effect in the data.^35, 36^ Instead, we aimed to directly quantify the probabilistic association of dimensions of traditional ADRD aetiopathology to social isolation to the extent supported by the observed data, while providing coherent estimates of associated uncertainty.

To this end, our analyses aimed at probabilistic answers to the question ‘How certain are we that loneliness/lack of social support is linked to an ADRD risk phenotype?’ Our analyses did not ask ‘Is there a strict categorical answer as to whether or not a risk phenotype is linked to loneliness or lack of social support?’ In this way, we aimed to directly quantify the population uncertainty intervals of risk effects in the context of social isolation. The full Bayesian model specification took the following form:

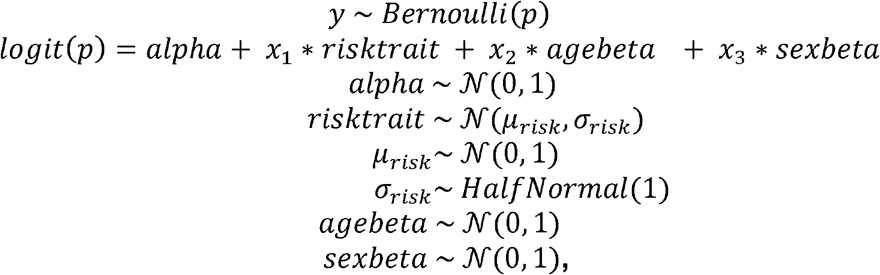

where *x*_1_ denotes an ADRD risk phenotype of interest (e.g., number of smoked cigarettes per day) and *y* denotes one of the target measures of social isolation (i.e., loneliness or lack of social support, cf. above). Details on the full list of examined risk traits can be in Supplementary Table 1. The multilevel formulation of the risk trait parameter serves flexible adaptation to different data settings. Variation that could be explained by participant age or sex was accounted for as potential confounds by the nuisance variables *x*_2_ and *x*_3_, respectively. In other words, in the UK Biobank or CLSA cohort, for a given risk phenotype of ADRD, we have estimated separate Bayesian models for loneliness and lack of social support.

Approximate posterior integration was achieved by means of Markov Chain Monte Carlo (MCMC), which sampled in a random walk towards the joint posterior distribution of all quantities at play.^32^ In 1,000 draws, the approximate parameter distributions were improved at each step in the sense of converging to the target distribution. At each step of the MCMC chain, the entire set of parameter values were estimated to be jointly credible given the data. In the data exploration phase, we have inspected model convergence by overlap between the geometry of posterior parameter distributions from four independent MCMC chains. We obtained further evidence for proper convergence to a stable model solution based on the effect sample size and 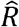 quality criteria. In the model exploitation phase, the final solution was computed by a single MCMC chain.

### Scientific computing implementation

Probabilistic hierarchical modeling and MCMC sampling^37^ were implemented as symbolic computation graphs in the *PyMC3* framework (https://github.com/pymc-devs/pymc3). Posterior parameter distribution plots were generated by *Seaborn* (https://seaborn.pydata.org/). Missing data was imputed using a nonparametric method for the UK Biobank, and a Bayesian method for the CLSA. All analysis scripts that reproduce the results of the present study are readily accessible to and open for reuse by the reader (http://github.com/banilo/to_be_added_later).

## Results

We set out to systematically explore possible links between major ADRD risk factors and rarely considered determinants of social isolation. Using a fully probabilistic approach, we carefully estimated the degree to which subjective and objective social isolation show population associations with established ADRD risk factors in the wider society. All our analyses reported in the following have been accounted for variation that can be explained by differences in participant age and sex. In the following, we present a series analyses of ADRD risk factors in four categories: 1) personal habits & lifestyle factors, 2) physical health factors, 3) mental health factors, and 4) societal & external factors, in parallel in similar measurements in the UKBB and the CLSA cohorts.

### Several rich cross-associations identified between social lifestyle and ADRD risk factors

We first performed a partial least squares analysis resulting in pairs of canonical vectors. The multivariate pattern-learning approach revealed the constellations of features that carry consistent associations within both high-dimensional variable sets, each comprising many features. The total variance explained of the original data matrices, shown separately for risk traits and social measures in Figure 1, is mapped for 7 PLS modes in the UKBB and 6 PLS modes in the CLSA. The canonical correlation for each mode quantified the linear correspondence between the two variable sets based on Pearson’s correlation between their canonical variates^38^. In both cohorts, a majority of the risk factors were linked to social lifestyle factors in at least one of the uncovered modes of joint variation.

**Figure 1:**
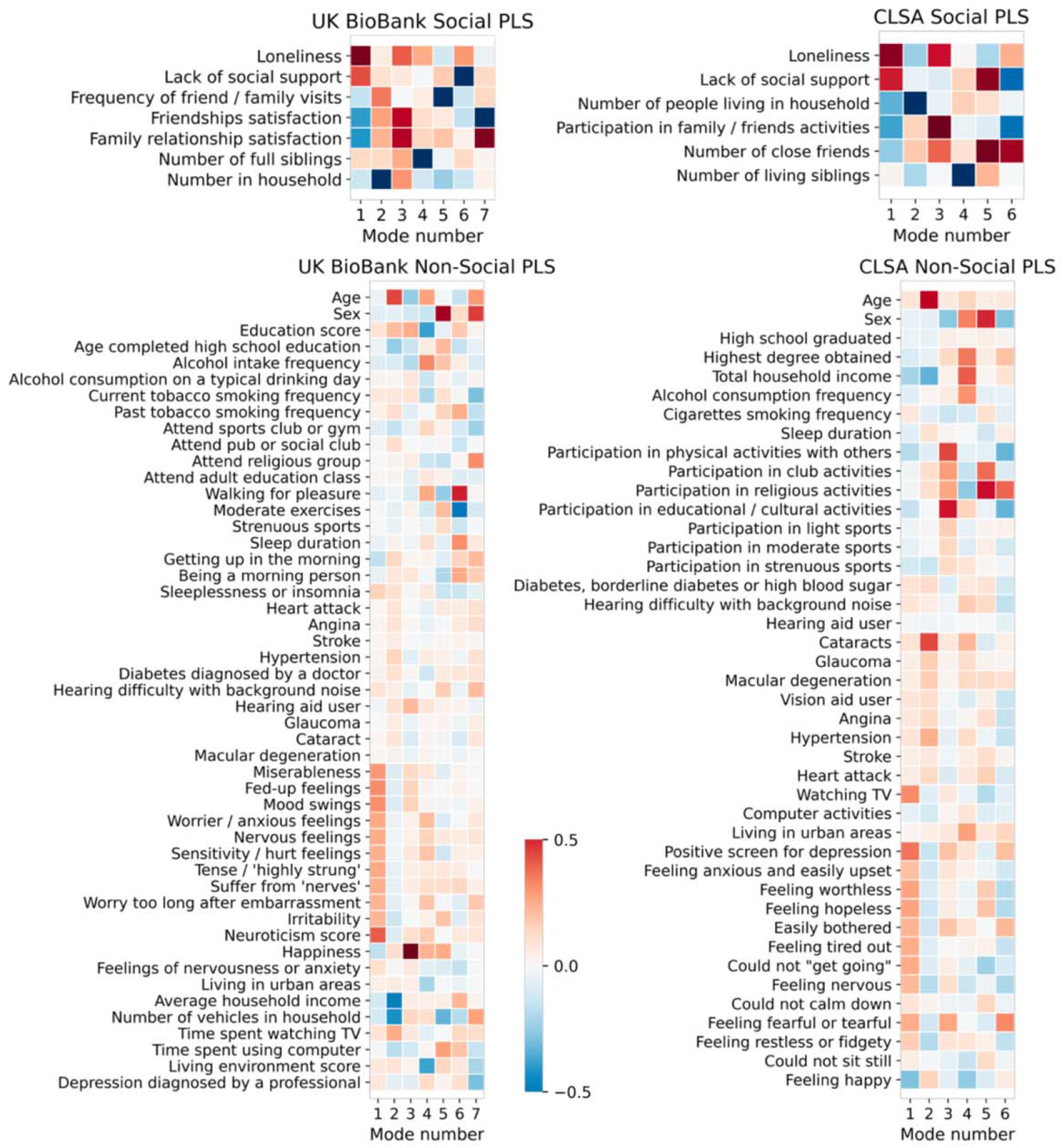
Widespread cross-associations exist between social deprivation indicators and traditional risk factors for Alzheimer’s dementias. To gain a synoptic overview, we initially explored multivariable relationships between sets of social isolation measures (top row) and sets of usually studied aetiopathological risk factors (lower row). In 502,506 UK Biobank participants (left column), the leading explanatory patterns (‘modes’) show that perceived and objective social isolation is associated with higher neuroticism scores and similar personality styles. In 30,097 CLSA participants (right column), the dominant pattern links loneliness and lacking social support to TV consumption and depression-related emotional traits. This doubly multivariate decomposition of two variable sets was obtained from partial least squares analysis (PLS; cf. methods). Note that this cursory analysis does not attempt to single out special variables (in contrast to the analyses from Figures 2–5). Overall, this perspective makes apparent that the majority of examined risk factors may be relate to some aspect of social lifestyle.

In the UKBB cohort, the first mode, by construction, explained a larger fraction of variation than any other mode, with a canonical correlation *rho* of 0.471 between the sets of variables. For the first canonical mode, interindividual differences in social richness dominated by loneliness (0.682) and lack of social support (0.437) were strongly paired with the personality traits among the ADRD risk factors, and the neuroticism score (0.408) in particular. Across the 7 modes, social determinants were related to lifestyle factors (e.g., exercise), physical health factors (e.g., hearing aid), mental health factors (e.g., personality), and societal factors (e.g., income). In the CLSA cohort, the variance in the first mode (*rho*=0.500) was best explained by interindividual differences in loneliness (0.652) and lack of social support (0.512) among the social factors, and by watching TV (0.321) and getting a positive screen for depression (0.364) for the risk traits. The PLS analysis on the UKBB and the CLSA indicated that the examined social determinants reflected the risk factors of ADRD from each of the four pillars in at least one of the uncovered modes of joint variation. Across both cohorts, the social domain of the first mode – which by construction, explains a larger fraction of variation than any other mode – was dominated by loneliness and lack of social support, which happen to be the two representative measures of subjective and objective social isolation throughout the present paper

### Bayesian regression between two determinants of social isolation and ADRD risk factors

Using a fully probabilistic approach, we next carefully estimated the degree to which subjective and objective social isolation show population associations with established ADRD risk factors in the wider society. Variation that could be explained by differences in participant age and sex was accounted for in all our reported analyses. The elected model was the same for all considered target risk factors. The full posterior parameter distributions – not sampling distributions – from our Bayesian modeling solutions for each variable of interest can be found in the Supplemental Material (cf. Supplementary Figures 1-4). For brevity, we here report the mean and the 90% highest posterior density interval (HPDI) of the model parameters, after seeing the data, which contains the 90% most credible parameter solutions in Table 1, summarized in the boxplots of Figures 2 to 5. The height of each boxplot refers to the mean value and the black error bars indicate the 90% HDPI of the effects of loneliness and lack of social support.

**Figure 2:**
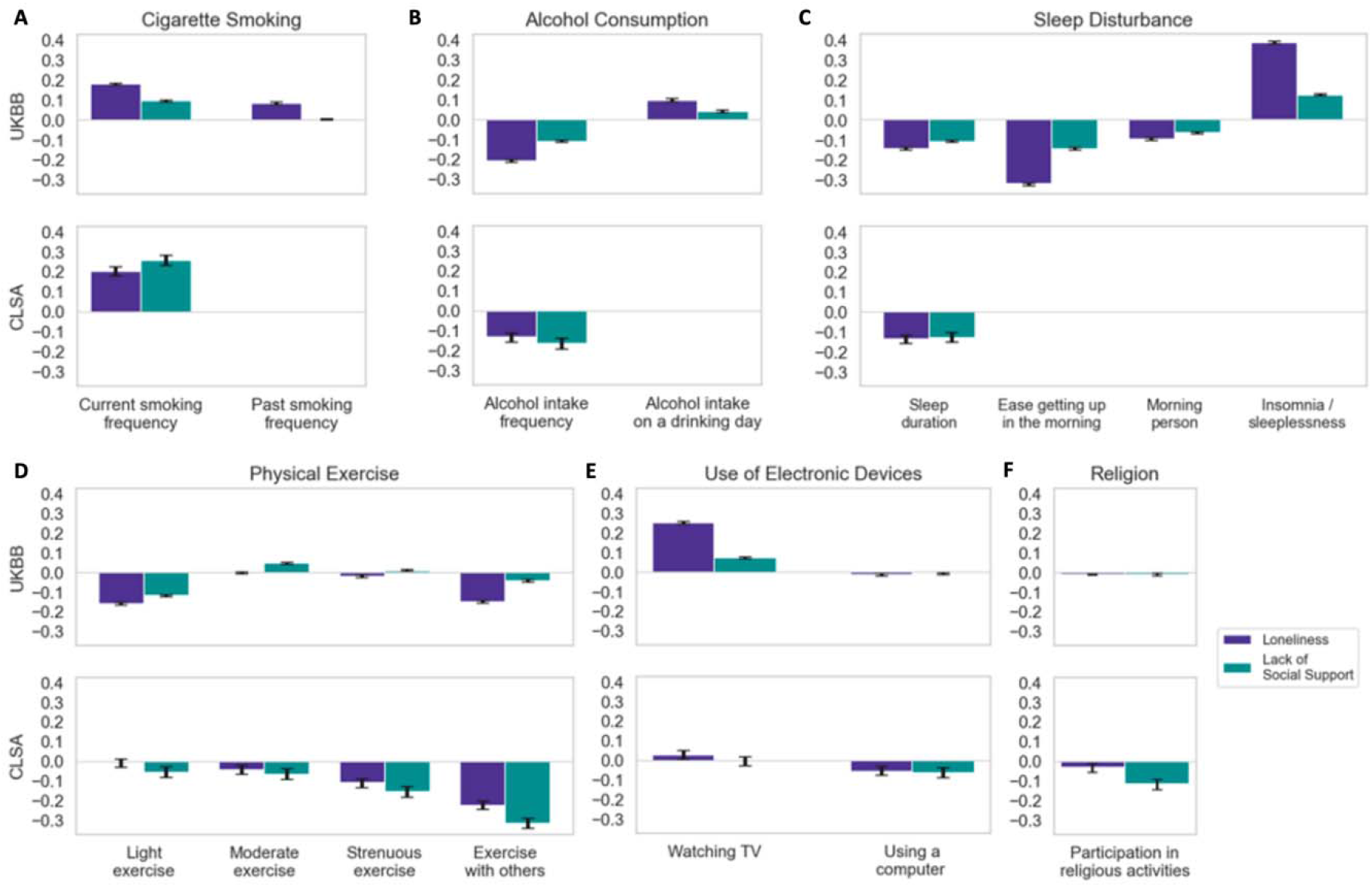
Various ADRD-related life style factors show strong association effects with loneliness and lack of social support across both cohorts. Bayesian estimation of the posterior probability that a given risk factor relates to one of two measures of social deprivation: loneliness and lack of social support. For simplicity, results are expressed as the mean and the 90% highest posterior density interval of the model coefficients (black error bars). In both UKBB and the CLSA, loneliness and lack of social support are robustly associated with a variety of lifestyle factors, including (A) current cigarette smoking, (B) alcohol consumption, (C) sleep duration, and (D) participation in physical activities with others. (E) Use of electronic devices and (F) participation in religious activities show smaller links to loneliness and weak social support. Both subjective and objective social isolation follow similar patterns in their associations with behavioural traits across the two cohorts. Sleeplessness has the largest effect association with social isolation in this category.

**Figure 3:**
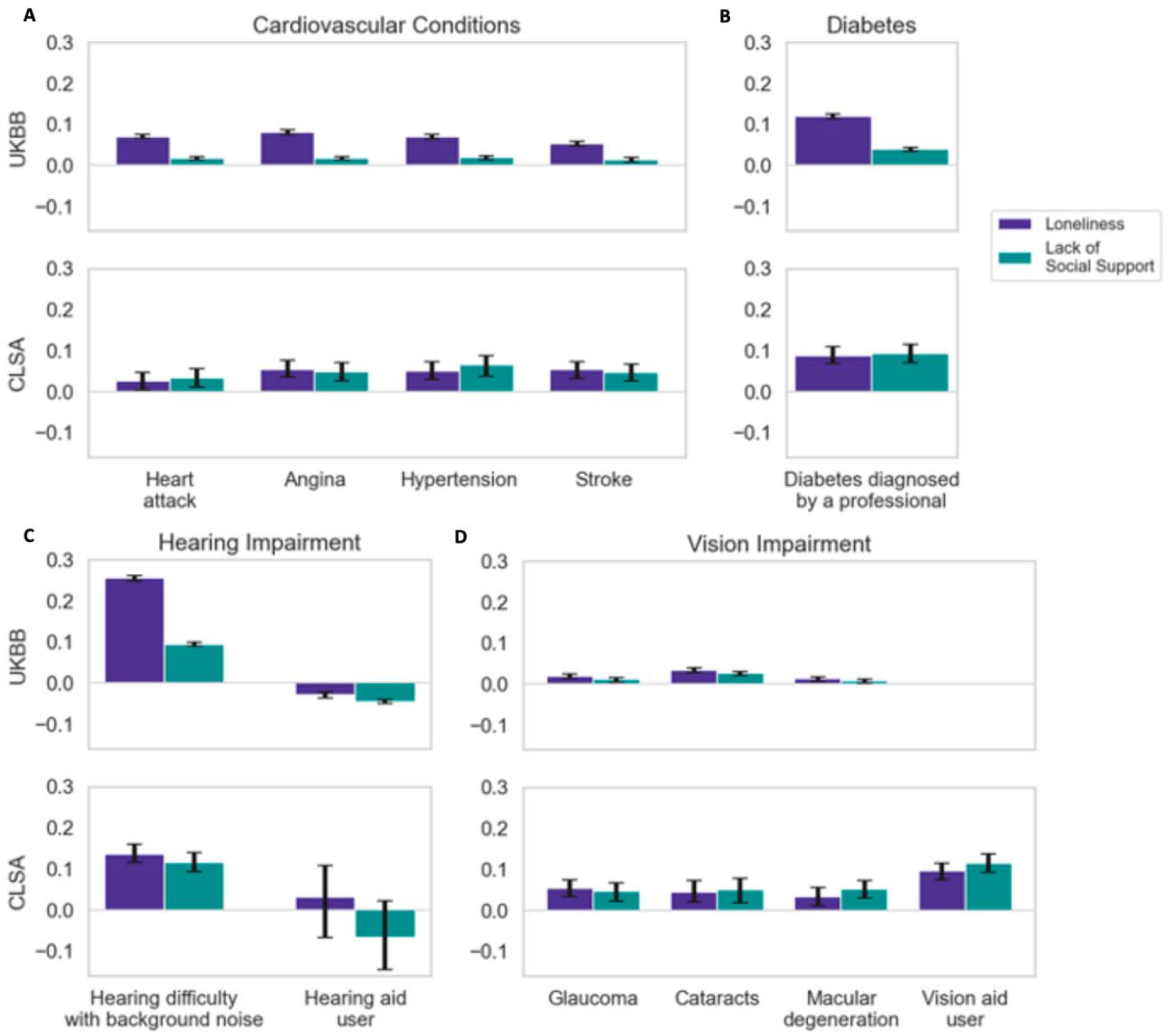
Physical health factors are related to social isolation. Bayesian estimation of the posterior probability that a given risk factor relates to one of two measures of social deprivation: loneliness and lack of social support. For simplicity, results are expressed as the mean and the 90% highest posterior density interval of the model coefficients (black error bars). In the UKBB and the CLSA cohorts, loneliness and poor social support show strong links with several physical health factors, such as (A) hypertension, (B) diabetes, (C) hearing difficulty with background noise, and (D) being a vision aid user. Across the two cohorts, hearing difficulty with background noise has the largest effect association with both subjective and objective social isolation in this pillar of risk traits.

**Figure 4:**
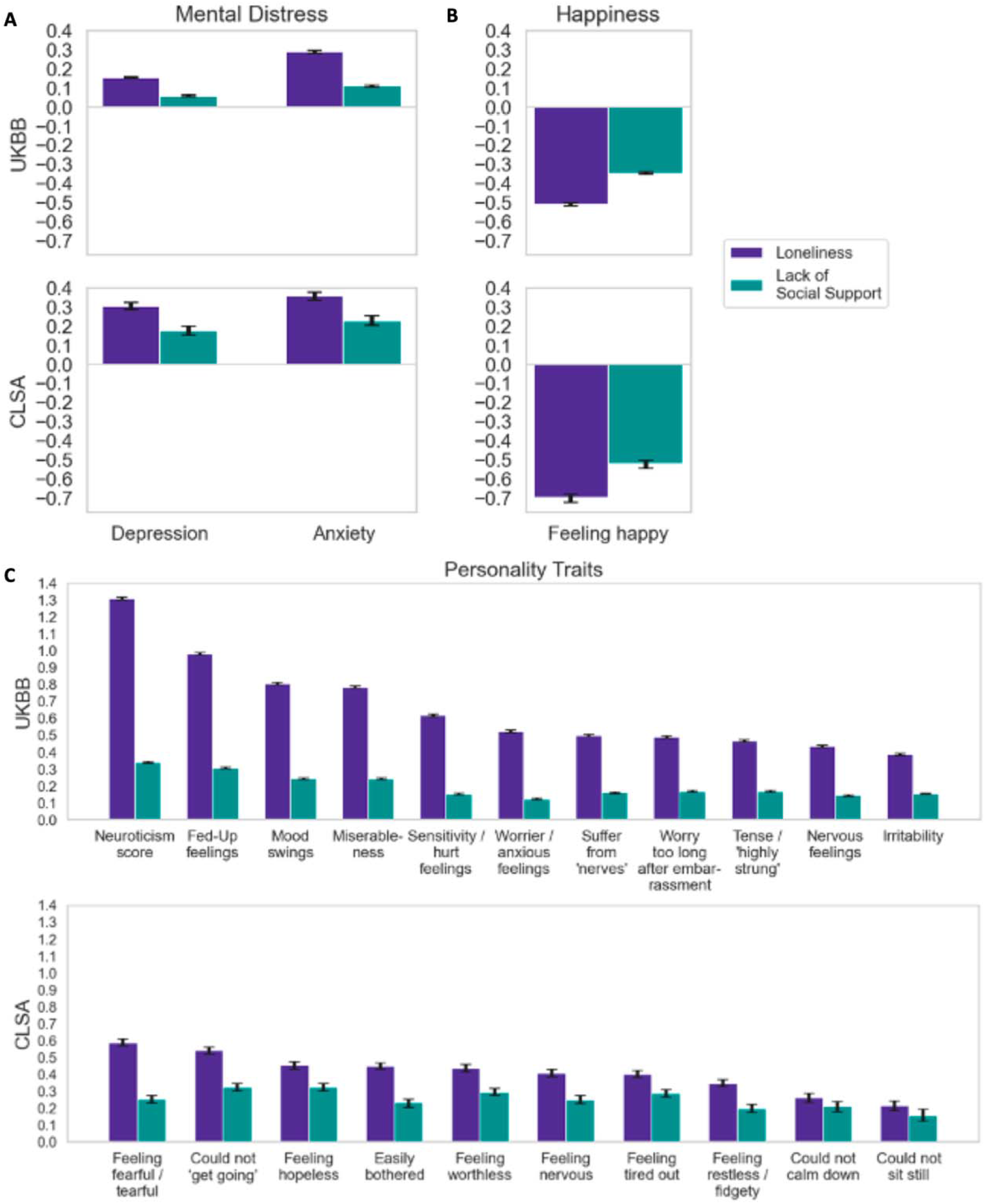
Mental health factors show prominent association effects with social isolation. Bayesian estimation of the posterior probability that a given risk factor relates to one of two measures of social deprivation: loneliness and lack of social support. For simplicity, results are expressed as the mean and the 90% highest posterior density interval for the model coefficients (black error bars). In this category and across the two datasets, both loneliness and lack of social support show some of the most prominent links with (A) depression and anxiety, (B) feelings of happiness, and (C) several measures of personality that play into the stress-buffer capacity of an individual. In particular, the neuroticism score has the largest effect association with both subjective and objective social isolation among all the examined ADRD risk factors.

**Figure 5:**
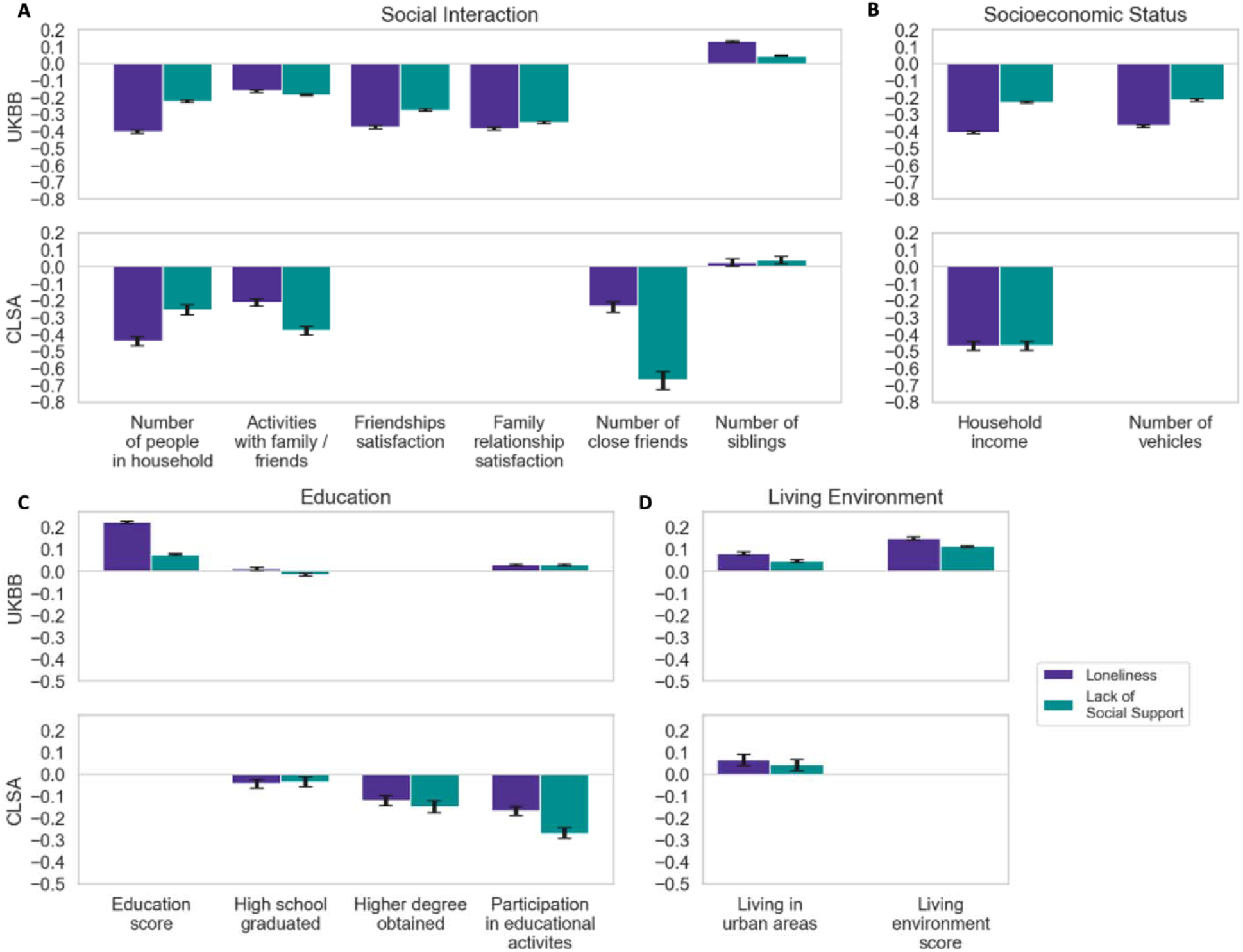
Societal risk factors exhibit salient association effects with social isolation. Bayesian estimation of the posterior probability that a given risk factor relates to one of two measures of social deprivation: loneliness and lack of social support. For simplicity, results are expressed as the mean and the 90% highest posterior density interval for the model coefficients (black error bars). In the UKBB and the CLSA datasets, loneliness and lack of social support show strong associations with several societal factors, including (A) the number of people living the household and the number of close friends, (B) household income, and (C) graduating from high school and obtaining higher degrees. (D) Living in an urban environment is also negatively linked with both measures of social deprivation.

**Table 1:**
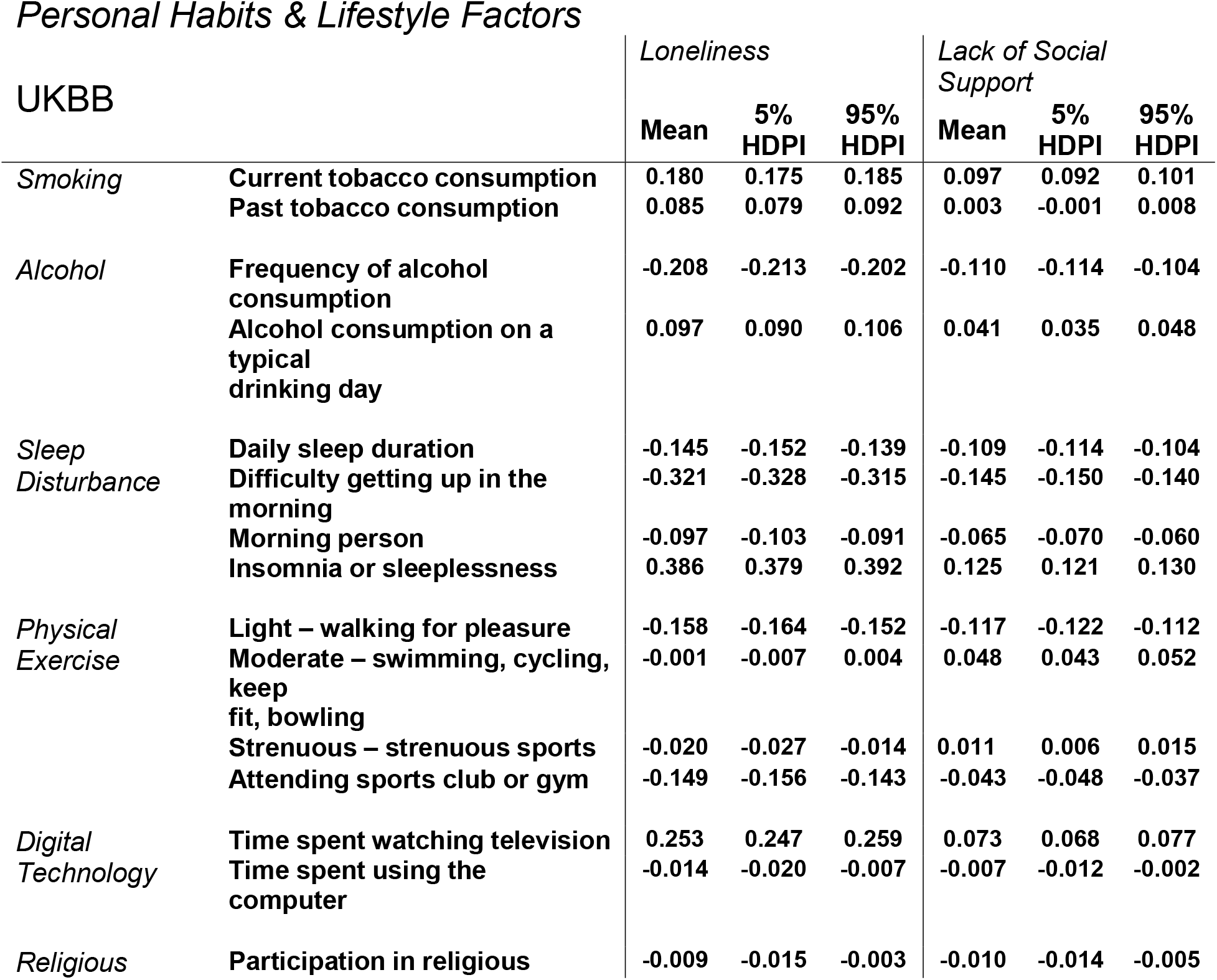
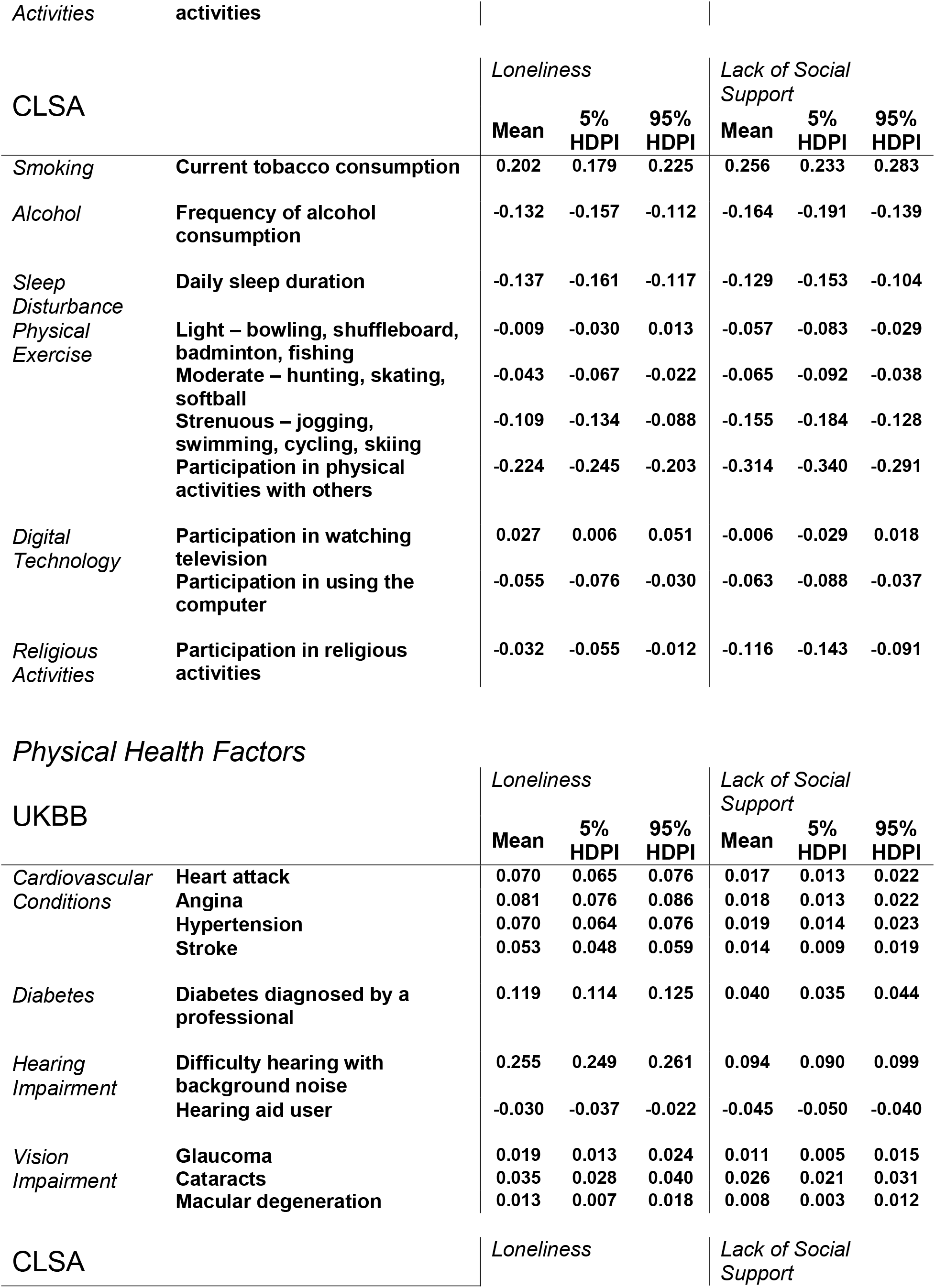
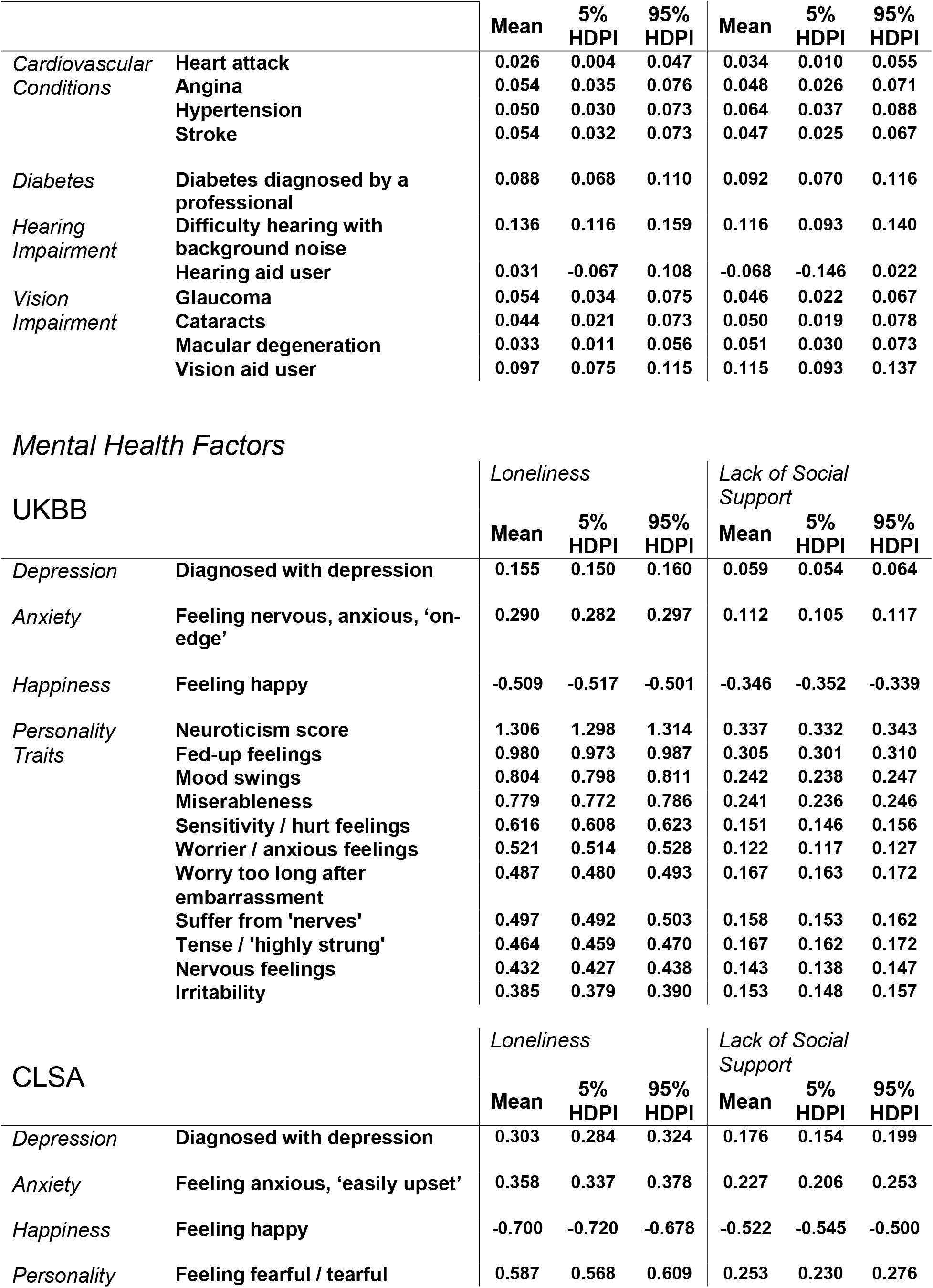
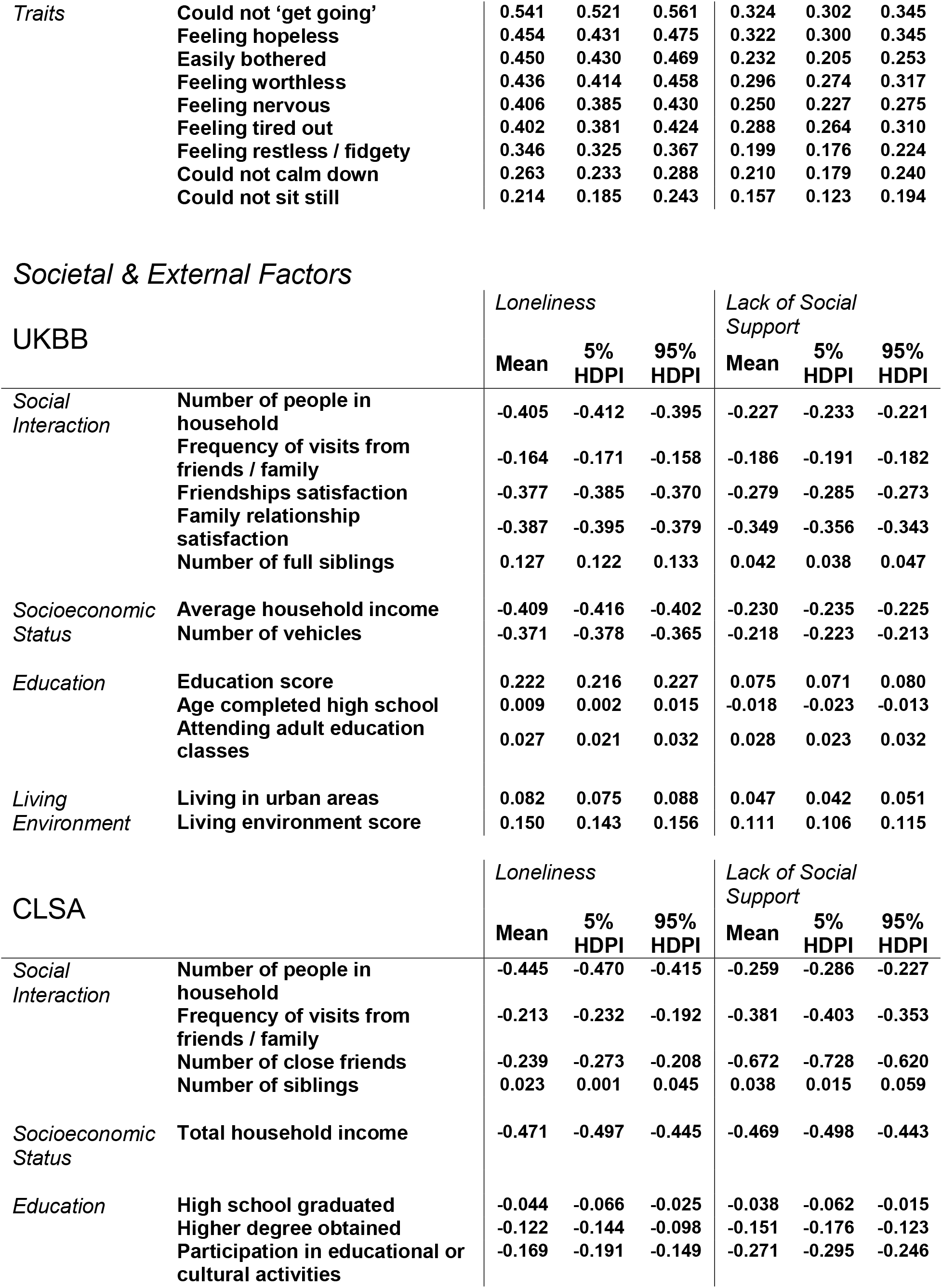
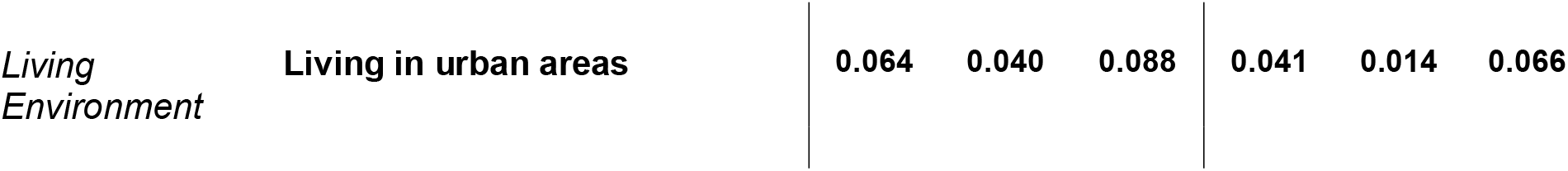
Mean associations between subjective and objective social isolation and ADRD risk factors in the UKBB and the CLSA.

#### Personal Habits & Lifestyle Factors

Taken together, our results showed statistically defensible links between both social determinants and the classical lifestyle risk factors of ADRD, which were replicated in the 502,506 UKBB and the 30,097 CLSA participants (Figure 2). Individuals who smoked more, excessively drank alcohol, experienced sleep disturbances, and failed to frequently participate in light-to vigorous physical activities were more likely to be lonely and lack social support. In the UKBB, for one unit increase in the number of cigarettes currently smoked, feeling lonely was 19.7% more likely than not feeling lonely. In addition, more frequent tobacco smoking corresponded to 10.2% increase in the odds of weak social support. In the CLSA, increasing regular participation in physical exercise with other people resulted in 20.1% decrease in the odds of feeling lonely and 26.9% decrease in having poor social support. Watching TV showed positive loneliness and weak social support effects. Instead, using the computer showed negative links with loneliness and poor social support. We also found negative effects from participating in religious activities on subjective and objective social isolation.

#### Physical Health Factors

We observed mutually confirmatory results between the CLSA and the UKBB among the physical health factors, with greater positive effect sizes on the social determinants found in the CLSA for cardiovascular diseases and visions impairments (Figure 3). Moreover, we discovered that vision-aid users in the CLSA were more likely to feel lonely and lack social support. Diabetes and hearing impairment, both recognized risk factors of dementia, showed prominent positive links with subjective and objective social isolation across both cohorts. In the UKBB, a unit increase in the difficulty to hear with background noise corresponded to a 29.0% increase in the odds of feeling lonely and a 9.86% increase in the likelihood of lacking social support. Still within the UKBB, individuals who used a hearing aid were less likely to be lonely and had weak social support. For the physical health factors in particular, the model uncertainty of our effects – indicated by wider posterior parameter distribution – was greater in the CLSA, attributable to the smaller set of data available in the CLSA relative to the UKBB.

#### Mental Health Factors

Collectively, mental health factors revealed strong population associations with both subjective and objective social isolation (Figure 4). All the different measures of personality, corresponding to neurotic and depressive behaviours, showed the largest positive associations with both subjective and objective social determinants, across the UKBB and the CLSA cohorts. In particular, the neuroticism score in the UKBB showed the greatest effect for loneliness and lack of social support. A greater neuroticism score accounts for 269.1% increase in the odds of feeling lonely and 40.1% in the odds of lacking social support. Further, we observed in both cohorts that feelings of happiness had a strong negative relationship with loneliness and poor social support. We also found relevant associations between an individual’s social capital and determinants of mental distress such as depression and anxiety.

#### Societal & External Factors

Overall, our results revealed that the opportunities for social interactions and the quality of these social exchanges held strong effect associations with loneliness and lack of social support in both datasets (Figure 5). In both cohorts, we found that individuals who shared their home with many people, and frequently participated in family or friendship activities were less often lonely and had better social support. In the UKBB, individuals who expressed greater satisfaction with their family relationship and their friendships revealed that the quality of social exchange also held salient effects on loneliness and lacking social support. And in the CLSA, a one unit increase in the number of close friends corresponded to 21.3% decrease in the odds of feeling lonely, and 48.8% decrease in the odds of lacking social support. However, in both the UKBB and the CLSA cohorts, we observed that having a greater number of siblings showed positive effects on loneliness and lack of social support. Further, we found salient links between the two measures of social isolation and socioeconomic status, measured as a combination of income, occupation, and education. In the UKBB, a unit increase in household income corresponds to a decrease in the odds of feeling lonely and lacking social support by 33.5% and 20.6%, respectively. Finally, in both the UKBB and CLSA, living in an urban environment, as opposed to a rural setting, showed a positive association with loneliness and poor social support.

## Discussion

The present study brings into sharp focus the multifaceted nature of interrelationships between social isolation and major ADRD risk factors. Our collective findings suggest that both perceived and factual social capital – loneliness and lack of social support – are consistently associated with classical ADRD risk factors, after accommodating effects for age and sex differences. To the best of our knowledge, this is the first study that explicitly targeted possible links between social isolation and a comprehensive array of most studied risk factors of ADRD, which we have here demonstrated using data from two nationally representative population cohorts of older adults from two different countries.

Among the examined measures of personal habits and lifestyle factors, sleep serves as a prototypical representative that showed several, and some of the largest, associations with social isolation, which successfully replicated across the UK Biobank and the CLSA cohorts. We found that all our measures of sleep disturbance had strong associations with loneliness and lack of social support across both cohorts. Similar to our findings, objective social isolation and self-reported loneliness have previously been linked to reduced sleep efficiency and poor sleep quality.^39–42^ Other investigators have hypothesized that perceived social isolation relates to hypervigilance for social threats,^20^ which in turn increases anxiety and reduces sleep quality. Consistent with this idea, many reports have shown that feelings of loneliness and reduced social support occur especially in individuals who report higher stress levels.^43–45^ Stress pile-up and emotional coping have been argued previously to contribute to the underlying reasons why lonely people are more often smokers,^46^ binge drinkers,^47^ and binge-watchers.^48, 49^ Interpersonal buffering, such as provided by subjective and objective social support, have been argued to be important psychosocial resources to cope with stressors.^46, 50^ There is a growing body of evidence that sleep disturbance,^51^ smoking cigarettes,^52, 53^ excessive alcohol consumption,^54^ and excessive television viewing^55, 56^ are all linked to cognitive decline and the development of ADRD. Our findings across two large cohorts showed that these potentially modifiable lifestyle factors that affect the onset of dementia have large effect associations with both loneliness and lack of social support.

Charting a series of physical health factors, notably cardiovascular conditions, diabetes, and physical exercise, we have shown all to feature some link to social isolation status, which corroborated across both cohorts. Our results are in line with existing research showing a detrimental effect of objective social isolation on subsequent dementia through cardiovascular pathways, by increasing the risk of hypertension^57^ and coronary heart disease.^58^ There is accumulating evidence that associates heart disease risk factors – diabetes, hypertension, obesity, smoking cigarettes – with late-life risk of cognitive impairment and dementia.^59, 60^ Moreover, participation in physical activities has been associated with better vascular health and lower risks of high cholesterol and diabetes,^61^ and regular physical exercise has repeatedly been shown to significantly reduce the risk of developing of dementia. Aside from the cardiovascular benefits, we explicitly showed that the social aspect of physical exercise was also important in relation to loneliness and social support. Consistent with our results, one study found that aerobic physical activities done alone did not seem to have any cognitive benefits in 754 healthy older adults.^62^

Among the considered mental health factors and all our examined measures in general, we found personality traits to feature the largest effect associations with social isolation, replicated across both cohorts. Our previous research has also shown a relation between personality traits and social isolation in genome-wide assessments in the UK Biobank.^12^ The neuroticism score, which reflects a person’s level of emotional volatility and vulnerability to stress, showed one the strongest effect size for loneliness and lack of social support, in the context of all the examined ADRD risk factors. Greater levels of late-life neuroticism have been previously associated with higher risk of developing mild cognitive impairment^63^ and dementia.^64–66^ Hostinar & Gunnar^67^ showed that through a phenomenon termed the social buffering of stress, specific personality traits can affect an individual’s susceptibility to the effects of stressors, while social support can dampen physiological stress responses.^68^

By the same token, the well-established ‘cognitive reserve’ hypothesis claims that intellectual enrichment provides a cognitive buffer to deal with injuries to the nervous system,^69^ which is an overarching theme among the societal factors In our rich datasets, we had the opportunity to concurrently examine the possible associations of numerous factors related to cognitive load, including education levels, socioeconomic status, computer use, sensory impairment, and different aspects of social interaction. Although, the relationship between social isolation and cognitive reserve is only now receiving increasing attention,^70–72^ we found consistently striking associations between these potentially modifiable societal factors and both loneliness and lack of social support, paralleled across two large cohorts, despite the slight difference in measures examined for the same construct between the two cohorts. Other investigators have suggested that interventions targeting social isolation and promoting a socially active lifestyle in later life may enhance cognitive reserve and reduce the risk of dementia.^71^

Given the current conceptual framework of this paper, our discussion revolves around dementia more broadly rather than Alzheimer’s disease in particular. In line with recent research on different biomarkers combinations in individual ADRD prognosis,^73^ our results also open the possibility for individual differences in the combinations of ADRD risk markers that are impacted by either or both subjective and objective social isolation. As our main contribution to the dementia literature, our study offers a comprehensive overview of the wide-ranging population-level associations between social deprivation and many ADRD risk factors.

## Conclusion

Our understanding about the implications of social isolation on ADRD remains in its infancy relative to the current evidence on other classical risk factors. However, our findings show a large array of associations between these potentially modifiable risk factors and both loneliness and lack of social support. Our collective findings underscore the importance of exploring subjective and objective social isolation in depth to inform policy interventions, especially among the elderly. Compared to other ADRD risk factors, such as ApoE4 genotype, social isolation is arguably easier to modify, and therefore, particularly promising to target and alter. As the persistence of the COVID-19 pandemic continues to force imposition of social distancing measures, research on these complex multiscale social phenomena may pave the way to address the two global public health priorities, separately recognized by the World Health Organization: ADRD and social isolation.

